# Optimizing genomic selection: A comparison of SNP selection strategies for reduced-density panels in beef cattle

**DOI:** 10.64898/2026.08.26.747408

**Authors:** Adebisi R. Ogunbawo, Henrique A. Mulim, Jorge Hidalgo, Henrique T. Ventura, Nadson O. Souza, Hinayah R. Oliveira

## Abstract

The exponential increase in the number of genotyped animals, combined with the availability of high-density SNP chips has introduced computational challenges for routine genomic evaluations, particularly during the construction of the genomic relationship matrix. Although higher-density SNP panels can facilitate the identification of causal mutations, their use substantially increases computational requirements without a proportional gain in genomic prediction performance. To optimize computational efficiency while maintaining accuracy of genomic predictions, this study compared five SNP selection strategies (i.e., random sampling, random sampling with inclusion of informative SNPs, linkage disequilibrium (LD)-based pruning, a Shannon entropy–based machine learning approach, and *F_ST_*-based prioritization) to develop reduced-density panels for Nellore cattle. Using high-density (HD) genotype data comprising 437,650 SNPs from 304,782 animals (after quality control) as reference, three reduced-density panels (25K, 45K, and 65K SNPs) panels were tested across five traits (i.e., Age at first calving, Stayability, Weaning weight, Yearling weight, Muscling) with diverse genetic architectures. Genomic estimated breeding values (GEBVs) derived from these reduced panels were compared to those obtained from the HD reference panel using Pearson’s correlations, under both genomic best linear unbiased prediction (GBLUP) and single-step GBLUP (ssGBLUP) methods. In the GBLUP model, prediction accuracy generally improved with increased marker density. Random selection with and without the informative SNPs consistently yielded the highest accuracies, whereas the *F_ST_*-based approach showed the lowest agreement with the HD reference across all densities. In contrast, ssGBLUP demonstrated strong robustness to marker reduction, producing uniformly high correlations (≈1.00) across all SNP densities and selection strategies. These findings indicate that optimized low-density SNP panels maintain prediction accuracy comparable to HD panels, offering a cost-effective tool for large-scale genomic evaluations.

**Author Summary:** Genomic selection has transformed cattle breeding by allowing producers to identify animals with superior genetic potential using DNA information. However, modern genomic evaluations often rely on very large genetic datasets that require substantial computing power and increase genotyping costs, particularly in large breeding populations such as Nellore cattle in Brazil. In this study, we evaluated whether reduced-density marker panels could maintain the same level of prediction accuracy as high-density panels commonly used in genomic evaluations. We compared five different strategies for selecting informative genetic markers and tested panels containing different numbers of markers across economically important traits. We found that reduced-density panels, particularly those developed using random or linkage-based selection methods, produced genomic predictions highly similar to those obtained with high-density panels. In addition, prediction methods that combined genomic and pedigree information remained highly robust even with fewer markers. Our findings suggest that reduced-density panels can support accurate and cost-effective genomic evaluations, allowing breeding programs to evaluate more animals more frequently while reducing computational demands.

## 1. Introduction

Traditionally, genetic evaluation in livestock species has primarily relied on phenotypic records and pedigree information to estimate breeding values [1–3]. The discovery of single-nucleotide polymorphisms (SNPs) marked a significant breakthrough in animal breeding, establishing genomic selection as a powerful tool for genetic improvement [1]. The integration of SNP data into genetic evaluations has significantly enhanced the accuracy of breeding value predictions [4,5] and accelerated the rate of genetic progress [6,7]. For instance, Wiggans et al. (2011) demonstrated that including genomic information in evaluations significantly improved the accuracy of breeding value predictions for Holstein, Jersey, and Brown Swiss bulls compared with traditional evaluations (traditional parent averages alone) [8]. Similarly, Kim et al. (2023) compared pedigree-based BLUP with single-step genomic best linear unbiased prediction (ssGBLUP) in Hanwoo cattle and found that including genomic information improved breeding value accuracy by 0.18 to 0.20 over the pedigree-only method [9].

Single-nucleotide polymorphisms are biallelic markers that occur abundantly across the genome [10]. Due to their dense coverage and association with quantitative trait loci (QTL), SNPs play a pivotal role in genomic prediction and the estimation of breeding values in livestock [11]. The SNPs are organized into commercial panels (or arrays) that enable efficient genotyping across populations [12,13]. The first medium-density SNP panel, the Illumina BovineSNP50 Bead Chip, was introduced in 2008 [14], followed by the high-density Bovine HD Bead Chip featuring approximately 777,000 SNPs [15]. Although these SNP panels have greatly facilitated the implementation of genomic selection in cattle, they were primarily developed using *Bos taurus taurus* reference populations [16]. As a result, these SNP panels have shown ascertainment bias when used in *Bos taurus indicus* breeds [16–18]. This bias may affect the accuracy and dispersion of genomic predictions in underrepresented breeds, as these panels often capture only a limited proportion of the genetic variance present in indicine (*Bos taurus indicus*) breeds [18].

The effectiveness of SNP panels in capturing the underlying genetic variation also depends on the extent of linkage disequilibrium (LD) between SNP markers and the causal variants influencing traits of interest [19,20]. In populations with strong and persistent LD, fewer SNPs may be sufficient to achieve accurate genomic predictions, as each marker can tag larger genomic segments. Conversely, in populations with weaker LD, higher SNP densities may be required to achieve comparable prediction accuracy [21,22]. Therefore, the informativeness of a SNP panel is not solely determined by the number of markers it contains, but also by how effectively those markers capture the LD structure of the target population. This interplay between SNP density and LD underscores the importance of tailoring genotyping strategies to the genetic architecture of specific breeds and production systems.

While increasing SNP density can improve prediction accuracy, several studies show this improvement reaches a plateau beyond a certain point [23–26]. Strategic reduction of SNP panel density could optimize computational resource utilization while maintaining prediction accuracy, enabling more frequent evaluations and broader population coverage in resource-constrained environments. This strategy becomes particularly relevant in resource-limited settings, where the rapid expansion in the number of genotyped animals places significant demands on existing computational infrastructure [27,28]. In developing countries, where server capacity is often limited, the burden of processing large-scale genomic data can hinder the routine implementation of genomic evaluations. As a result, many institutions opt for low- or medium-density SNP panels for genomic evaluations. For instance, the Brazilian Association of Zebu Breeders (ABCZ) currently uses a 120K SNP panel for genomic evaluations of Nellore cattle. However, currently, genomic evaluation reports are published only four times per year, limiting the frequency and responsiveness of genetic assessments.

One potential solution to this bottleneck is to reduce the SNP panel density, thereby lowering computational demands while preserving prediction accuracy relative to the currently used 120K panel. Moreover, a more efficient panel would likely allow more animals to be genotyped at lower cost and evaluated more frequently, ultimately enhancing the accuracy of genomic selection and accelerating genetic progress in the Nellore population. However, despite the growing interest in reduced-density panels, there is currently no consensus on the optimal strategy for selecting informative SNPs. Various *ad hoc* methods have been proposed, but few studies have directly compared their performance in terms of prediction accuracy, computational efficiency, and practical impact on genomic selection. Therefore, the objectives of this study were to: (1) compare five SNP selection strategies for developing reduced-density panels in Nellore cattle; and (2) evaluate the impact of these selection methods on genomic prediction accuracy by comparing breeding values obtained from GBLUP and ssGBLUP against those derived from high-density and 120K (ABCZ) reference panels.

## 2 Results

### 2.1. Comparison of SNP panels generated by different selection methods

The pairwise comparisons within the three SNP densities (25K, 45K, 65K) revealed substantial variability in marker overlap among selection scenarios, indicating that different strategies prioritize largely distinct genomic regions. As expected, the Random and Random including Significant scenarios were consistently the most similar scenarios used. In contrast, Random-based approaches showed limited overlap with LD, *F_ST_*, and Shannon entropy–based strategies, suggesting that these methods capture different patterns of genomic variation. Among the non-random approaches, LD and Shannon entropy consistently exhibited the greatest similarity across all densities, whereas *F_ST_* showed comparatively lower overlap with the other methods. Although the absolute number of shared markers increased with marker density, the relative relationships among selection strategies remained stable. The average proportion of shared SNP markers across different scenarios, considering the three SNP densities together, is shown in Table 1. The specific counts of SNP markers shared and unique across selection scenarios, for each SNP density, are available in S1a-c Tables.

**Table 1:**
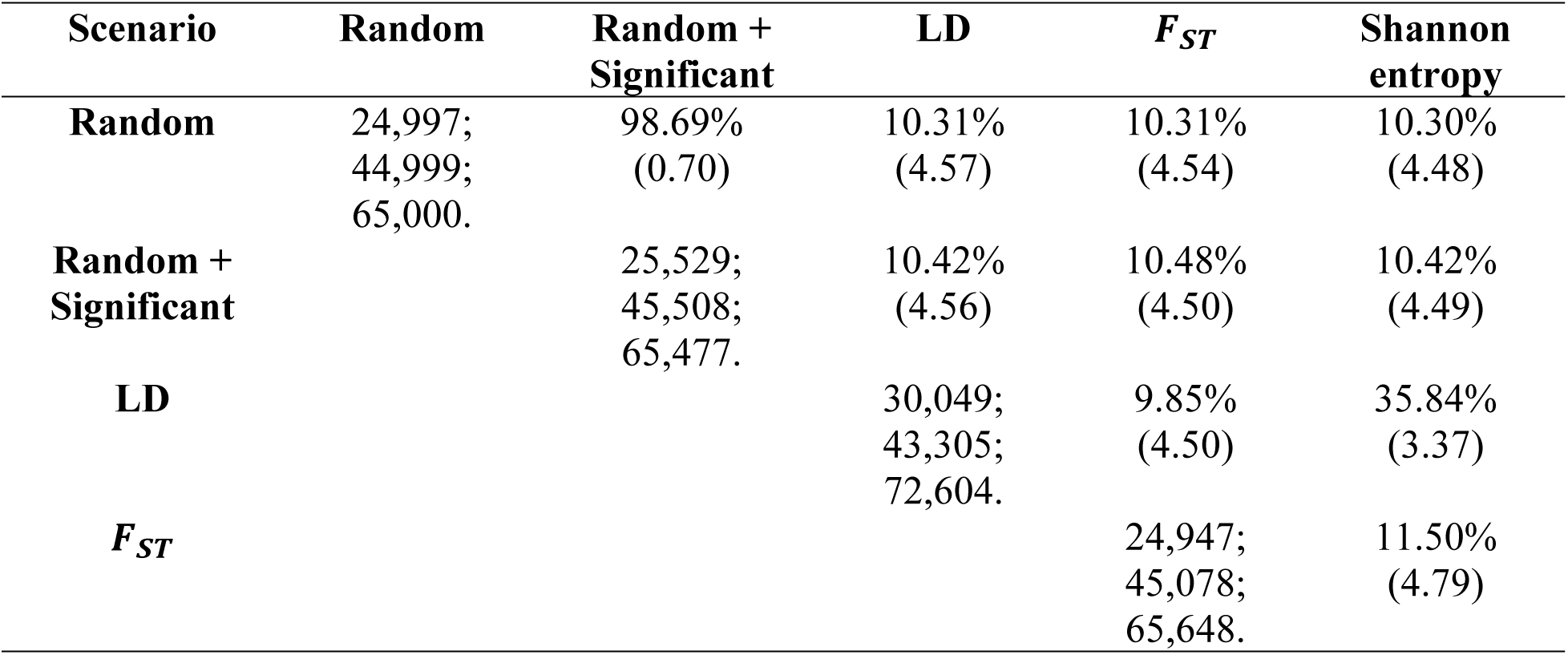
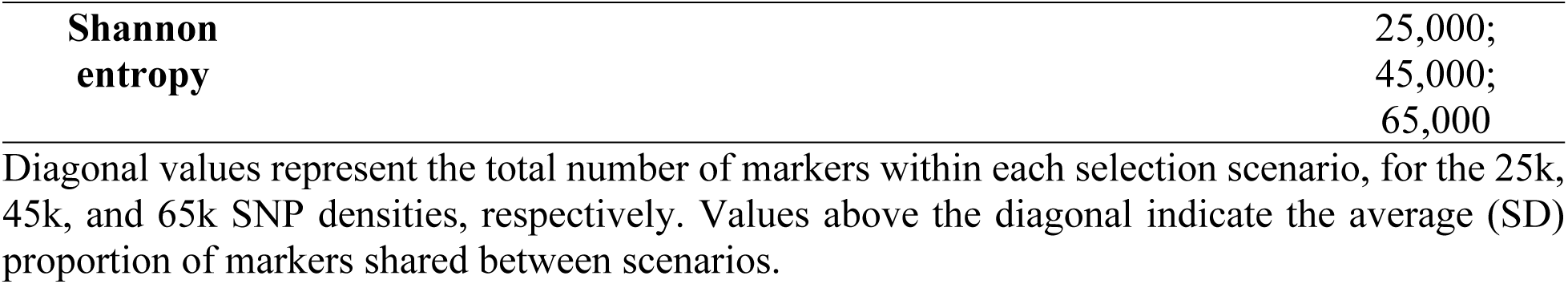
Average proportion of shared SNP markers across different scenarios. The tested scenarios include Random, Random with significant markers, LD, F_ST_, and Shannon Entropy. Proportions included in this table are averages (SD) of the proportions calculated for each SNP density tested (i.e., 25k, 45k, and 65k).

### 2.2 Accuracy of genomic predictions for the different selection scenarios across traits

To evaluate the performance of the five SNP selection strategies across the evaluated traits, Pearson correlation coefficients were computed between predicted GEBVs obtained from the HD reference panel and each combination of prediction method (GBLUP vs. ssGBLUP), selection scenario (Random, Random with significant markers, LD, F_ST_, and Shannon Entropy), and SNP density (25K, 45K, and 65K). The resulting prediction accuracies obtained for each trait are shown in Fig 1. Specific values are provided in the S2a-j Tables.

**Fig 1.**
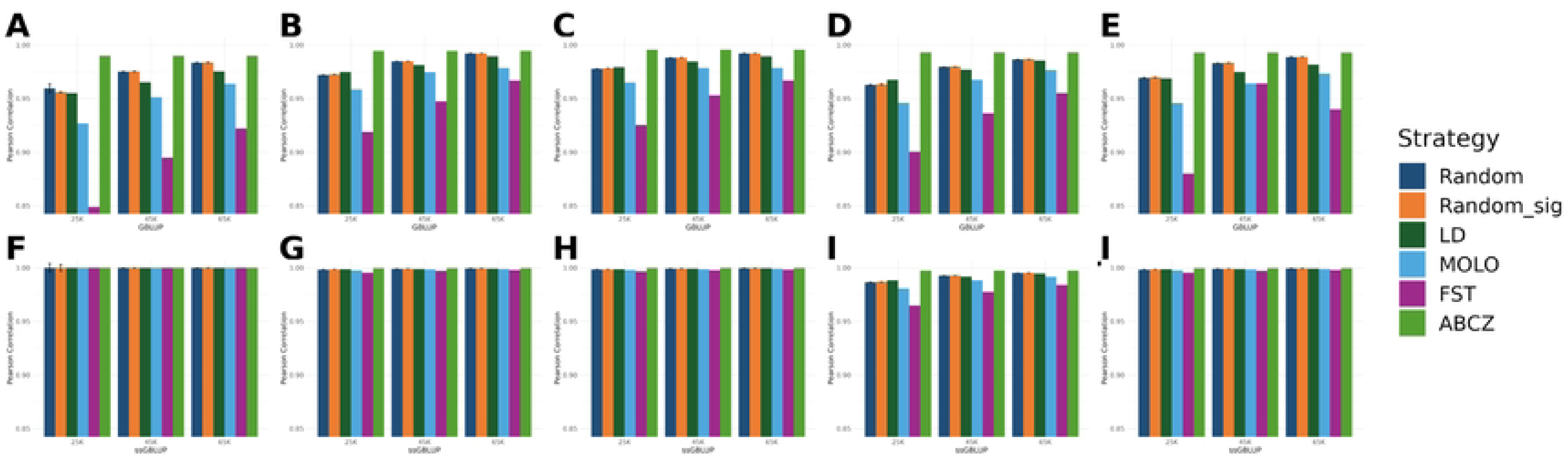
Accuracy of genomic predictions across traits, SNP densities, selection strategies, and prediction models. Panels A-E show results from the GBLUP method for age at first calving (AFC), weaning weight (WW), yearling weight (YW), Muscling score (MUSC) and stayability (STAY), respectively. Panels F - J show the corresponding results from the ssGBLUP method across the same traits.

Across all evaluated traits, consistent patterns in genomic prediction accuracy were observed across selection strategy, SNP density, and prediction methods. Under the GBLUP method, prediction accuracy increases consistently as marker density expanded from 25K to 65K across all SNP selection strategies (Fig 1). However, the magnitude of improvement diminished at higher densities, suggesting that moderate-density panels (e.g., 45K–65K SNPs) can effectively capture most genomic information required for accurate prediction. Among the selection strategies evaluated under the GBLUP framework, the Random, Random with inclusion of significant SNPs, and LD-based selection strategies showed similar performance and yielded the highest agreement with the HD reference panel for all traits. In contrast, the Shannon entropy and F_ST_-based selection approaches consistently showed the lowest agreement with the HD panel across traits and panel densities, suggesting that these approaches are less effective for optimizing genomic prediction in this population.

In contrast to GBLUP, prediction accuracies obtained under the ssGBLUP method were uniformly high and largely insensitive to both SNP density and selection strategy across traits (Fig 1). For instance, correlations between GEBVs derived from the reduced-density panels and the HD reference approached unity (∼1.0) across all traits and scenarios. This near-perfect agreement demonstrates that ssGBLUP is highly robust to marker reduction, likely because integrating comprehensive pedigree and phenotypic data effectively compensates for the loss of marker information at lower densities. Finally, while the 120K SNP panel currently utilized by ABCZ exhibited the highest overall agreement with the HD reference across all traits and models, the reduced-density panels, particularly those developed via random and LD-based selection at 45K and 65K densities, achieved highly comparable accuracies. These results underscore the potential of strategically reduced panels as cost-effective and computationally efficient alternatives for routine genomic evaluations in Nellore cattle.

Strategy: (Random) markers randomly selected from HD panel; (Random_sig) inclusion of significant markers to random strategy, (LD) selection based on linkage disequilibrium pruning, (Shannon) selection based on Shannon entropy, (F_ST_) selection based on marker prioritization, and (ABCZ) the current SNP panel used by the Brazilian Association of Zebu breeders.

## 3. Discussion

Several selection strategies have been proposed to design SNP arrays or select SNPs from an HD SNP panel to develop low-density alternatives [29]. Low-density panels are still significantly cheaper, facilitating broader population coverage [13]. These SNP selection strategies aim to retain the most informative markers [13,29,30] while simultaneously reducing genotyping costs and computational demands. Despite the increasing use of reduced-density panels, there is still no clear consensus on the most effective strategy for selecting informative SNPs. Several *ad hoc* methods have been proposed [13,29,31], yet few studies have compared these strategies in terms of prediction accuracy, computational efficiency, and their overall impact on genomic selection. Consequently, this study assessed five distinct SNP selection strategies across varying SNP densities, prediction models, and traits with diverse genetic architectures to evaluate their ability to maintain prediction accuracy relative to the HD reference panel.

When comparing the different selection strategies, the proportion of overlapping SNPs was generally low (Table 1), with most markers being unique to each selection approach. Interestingly, the proportion of shared SNPs remained stable across the different panel densities, suggesting that each selection strategy consistently prioritizes distinct genomic features regardless of the total number of markers selected. Among all approaches, the Random and Random with inclusion of significant markers strategies showed the highest level of overlap, sharing approximately 98.69% of SNPs. This high concordance was expected, as both approaches rely on largely identical marker sets, with the distinction that the latter includes unique markers with known associations, effectively balancing broad genome coverage with the inclusion of informative loci. However, adding these significant SNPs to this set of random scenarios did not necessarily result in a substantial change in prediction accuracy across densities (Fig 1), likely because the number of significant markers included represented less than 1% of the total panel size (Table 1).

Pairwise comparisons among the other strategies (e.g., Random vs. LD, Random vs. *F_ST_*, Random vs. Shannon entropy, Random with inclusion of significant markers vs. LD, Random with inclusion of significant markers vs. *F_ST_*, Random with inclusion of significant markers vs. Shannon entropy, LD vs. *F_ST_*, and *F_ST_* vs. Shannon entropy) showed minimal overlap in the proportion of shared SNPs across densities, ranging from 9.85% to 11.50% (Table 1). The lowest overlap (i.e., 9.85%) was observed between the LD- and *F_ST_*-based strategies. These results indicate that the SNP sets selected by each strategy strongly reflect their distinct underlying prioritization criteria. A notable exception was the comparison between the LD-based and Shannon entropy strategies, which showed a substantially higher overlap across SNP densities (35.84%). This greater overlap likely reflects the tendency of both methods to prioritize highly informative SNPs and maximize the amount of genetic information captured per marker (Wu et al., 2016). Specifically, Shannon entropy selects SNPs with high allelic diversity and information content [13], while LD-based selection targets markers that effectively capture variation within haplotype blocks. Consequently, entropy-based methods frequently identify SNPs that also function as effective LD tag markers [32–34], leading to the observed overlap between these two scenarios.

Under the GBLUP method, prediction accuracy consistently increased as marker density expanded from 25K to 65K, relative to the HD panel. For instance, the Pearson correlation for AFC using random selection increased from 0.9596 at 25K to 0.9751 at 45K and reached 0.9836 at 65K (Fig 1 and S2a Table). Across all densities, the Random approach, Random with inclusion of significant markers, and the LD selection scenario had the highest correlation with the HD panel. The exceptionally high agreement observed for the random-based approaches is likely due to the stratified random sampling method used across chromosomes, where the number of SNPs per chromosome was proportional to its physical size. This stratification ensured that SNPs were evenly distributed across the genome, thereby maintaining adequate genome-wide representation and preserving the genomic relationship structure captured by the HD panel [35,36]. Similarly, the LD scenario exhibited high correlations with the HD panel because this approach prioritizes tag SNPs that represent haplotype blocks across the genome [32–34], which also effectively preserves genomic relationships among individuals.

Although the magnitude of improvement was modest, the random selection approach with inclusion of prioritized markers showed slightly higher correlations with the HD panel compared to the purely random selection scenario under the GBLUP method. This consistent trend suggests that integrating previously identified, trait-relevant markers can enhance the informativeness of reduced-density panels. Conversely, the *F_ST_* scenario performed poorly compared to the other selection strategies, although the accuracy for this selection strategy improved with increasing density (e.g., 0.9188, 0.9476, 0.9668 for WW at 25K, 45K, and 65K, respectively). This poor performance in the GBLUP method likely occurs because the *F_ST_* based selection in GBLUP models across most of the traits could be a result of the *F_ST_* approach, which measures genetic differentiation over time, which does not necessarily prioritize SNPs that are in high LD with causal variants for the traits evaluated [31]. The Shannon entropy scenario yielded intermediate results (i.e., lower correlation than the Random, Random with inclusion of significant markers, and LD-based scenario, but higher than the *F_ST_* scenario), with accuracy also increasing alongside marker density (Fig 1).

In contrast to GBLUP, when evaluating the ssGBLUP method [37–39] across the five traits, the Pearson correlations between GEBVs obtained from the reduced SNP densities (25K, 45K, and 65K) and the HD reference panel were approximately 1.00. These near-perfect correlations were consistent across all selection strategies, indicating that increasing marker density within this range did not meaningfully affect GEBV prediction. This stability suggests that the additional pedigree and phenotypic information incorporated in ssGBLUP makes this method highly robust and less sensitive to reductions in SNP density. Consequently, for populations with deep pedigree recording, a 25K SNP panel appears to be as informative as a 65K panel for GEBV prediction.

## 4. Conclusion

This study demonstrates that reduced-density SNP panels can achieve genomic prediction accuracy similar to that of high-density SNP panels in Nellore cattle. Although prediction accuracy under the GBLUP method increased with SNP density across all evaluated traits, the marginal gains diminished at higher densities, indicating that panels in the range of 45K–65K SNPs capture most of the genomic information required for reliable genomic evaluation. Furthermore, the ssGBLUP method proved highly robust to marker reduction, maintaining near-perfect prediction accuracy even at the lowest density evaluated (25K). Overall, these findings highlight that reduced-density panels developed using informed SNP-selection strategy (such as those developed using LD-based selection or random sampling inclusive of significant markers) may provide a cost-effective and computationally efficient alternative to high-density panels. Implementing these optimized panels can facilitate more frequent genomic evaluations, enable broader population coverage, and improve overall scalability of genomic selection programs, particularly in resource-constrained environments.

## 5. Material and Methods

Animal Care and Use Committee approval was not needed for this study, as all data were obtained from an existing database.

### 5.1 Original dataset and genotype imputation

The Brazilian Association of Zebu Breeders provided the variance components, pedigree, phenotypes, and genotypes used in this study. The pedigree information was available for 14,602,423 Nellore animals raised in Brazil, including 6,603,793 sires, 7,995,760 dams, and 4,095,918 alternate dams. A total of 304,782 animals were genotyped using 14 commercially available SNP panels, which ranged from low to high density. After defining the optimal imputation approach for this population [40], imputation was performed using the FImpute v3 software [41], considering pedigree information. Imputation was performed in two steps: first, all low- and medium-density genotypes were imputed to a custom SNP panel containing approximately 120K SNPs, created by combining the 50K and 70K panels (over 86K animals in the reference). Subsequently, the 120K SNP panel was imputed to the high-density SNP panel (777K), using 1,962 animals genotyped with the high-density SNP panel in the reference. Quality control performed before imputation retained only SNPs mapped to autosomes with a call rate above 0.90, minor allele frequencies above 0.01, and deviation from the Hardy-Weinberg equilibrium below 0.15. Imputation accuracy was greater than 0.98 for the dataset used. Additional details on imputation performance for this dataset can be found in Campos et al. (2026) [40].

The phenotypic dataset was available for five traits: age at first calving (AFC), stayability (STAY), weaning weight (WW), yearling weight (YW), and muscling score (MUSC). These traits were selected to represent different genetic architectures and biological pathways, ensuring that the developed SNP panels would be robust enough for all traits currently evaluated by ABCZ and adaptable to new traits that might be added in the future. For instance, AFC and STAY represent reproductive traits with moderate to low heritability (evaluated using a linear and threshold model, respectively), WW and YW represent growth traits with moderate heritability (linear models with and without maternal effects, respectively), and MUSC represents a subjectively scored trait with high heritability (evaluated using a linear model). Details about the phenotypic traits and how they are recorded by ABCZ can be found in Ogunbawo et al. (2025) [42].

### 5.2 Phenotypic and genotypic quality control

Before quality control, the dataset comprised 5,184,615 observations for AFC; 1,206,657 for STAY; 4,462,467 for WW; 3,778,244 for YW; and 237,248 for MUSC. Quality control was performed on the phenotypic records, with missing observations and phenotypic records exceeding +/- 3 standard deviations from the contemporary group (CG) average being removed for all traits except STAY. For STAY, the quality control only removed CGs without variation in the STAY phenotype. The CGs were created based on the information of herd-year-season of birth, herd-year-season of measurement, and day of measurement to account for the effect of technician/farmer (for all traits except STAY, which included only herd-year-season of birth in the CG). The CGs that had fewer than 3 animals were also removed to ensure data accuracy, reliability, and consistency. The total number of observations/animals remaining after quality control and descriptive statistics for each trait are shown in Table 2.

**Table 2.**
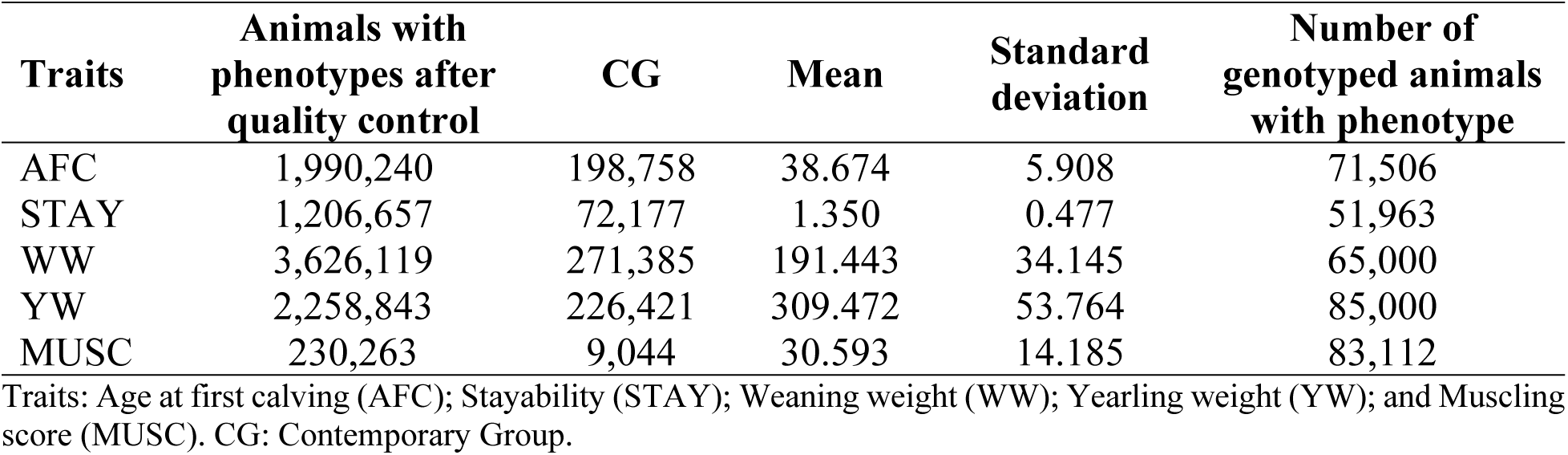
Descriptive statistics for age at first calving (AFC), stayability (STAY), weaning weight (WW), yearling weight (YW), and muscling score (MUSC), after quality control.

| Traits | Animals with phenotypes after quality control | CG | Mean | Standard deviation | Number of genotyped animals with phenotype |
| --- | --- | --- | --- | --- | --- |
| AFC | 1,990,240 | 198,758 | 38.674 | 5.908 | 71,506 |
| STAY | 1,206,657 | 72,177 | 1.350 | 0.477 | 51,963 |
| WW | 3,626,119 | 271,385 | 191.443 | 34.145 | 65,000 |
| YW | 2,258,843 | 226,421 | 309.472 | 53.764 | 85,000 |
| MUSC | 230,263 | 9,044 | 30.593 | 14.185 | 83,112 |

Traits: Age at first calving (AFC); Stayability (STAY); Weaning weight (WW); Yearling weight (YW); and Muscling score (MUSC). CG: Contemporary Group.

Genotypic quality control was performed using the QCF90 software, part of the BLUPF90 family programs [43]. Quality control was performed to exclude SNPs based on minor allele frequency (MAF) < 0.05, SNP call rate < 0.90, animal call rate < 0.90, extreme deviation from Hardy-Weinberg equilibrium (>0.15; calculated as the difference between observed and expected heterozygote frequencies; following [44], and SNPs located on non-autosomal regions. Finally, a total of 304,782 animals and 437,650 SNPs distributed across the 29 autosomal chromosomes remained for further analysis.

### 5.3 Development of SNP panels

Deciding on the appropriate number of SNPs required to maximize the accuracy of genomic prediction while minimizing SNP number to optimize evaluations is a crucial step in determining the optimal density of a SNP panel. Knowing the extent of LD in the bovine genome is one of the steps required to decide on the appropriate number of SNPs or markers that will be sufficient for genomic studies, as LD helps quantify the amount of information that can be inferred about one locus from another [20,45]. Recent estimates of LD decay in Nellore cattle indicate an average LD of 0.282 ± 0.271 and a mean distance between adjacent SNPs of 31,567.6 bp across all chromosomes [40]. Based on the ARS-UCD1.2 bovine genome assembly used in our study, with an approximate genome length of 2.71 Gb, a target marker spacing of 45 kb corresponds to approximately 60,222 evenly distributed SNPs [40]. Therefore, a panel of approximately 65,000 SNPs was selected as the upper reduced-density panel to provide a slightly greater marker coverage.

To ensure fair comparison among the different *ad hoc* methods tested and determine the optimal number of SNP markers required for accurate genomic selection, we selected three different SNP panel densities to test: approximately 25,000 (25K), 45,000 (45K), and 65,000 (65K). These SNP panels were selected from the high-density panel after quality control (N = 437,650 SNPs) to represent a range of low-density panels capable of capturing varying levels of LD across the Nellore genome. To provide a population-specific context for the SNP densities, the effective number of independent chromosome segment (Me) was approximated using PREGSF90 in BLUPF90+ suite [43] as the number of largest eigenvalues required to explain 98% of the variation in the genomic relationship matrix (G), as proposed by Pocrnic et al. (2016) and Misztal et al. (2020). This analysis resulted in an estimated genomic dimensionality of 36,021 independent chromosome segments [46–48]. Therefore, the 25K, 45K, and 65K panels were selected to span marker densities below and above the estimated genomic dimensionality (*M_e_* = 36,021) of the Nellore population, allowing evaluation of whether increasing SNP density beyond the estimated effective dimensionality resulted in additional gains in genomic prediction accuracy.

For each SNP density, five different methods for SNP selection were compared: random selection, random selection including significant markers from previous genome-wide association studies (GWAS), LD-based selection, *F_ST_* based selection [49,50], and Shannon entropy-based selection [29,51]. A scheme of all scenarios created from the HD SNP panel is shown in Fig 2.

**Fig 2.**
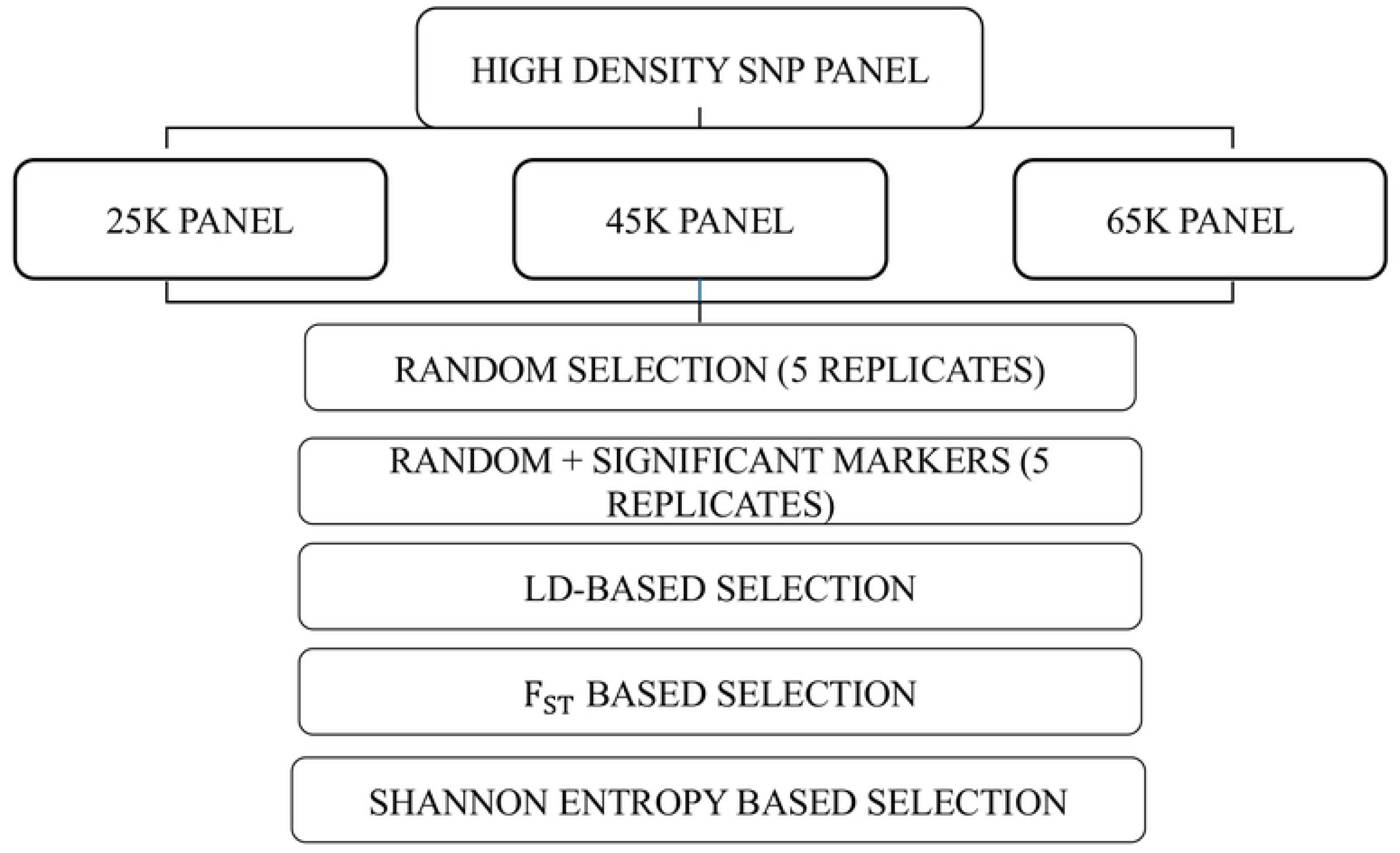
Scheme of SNP selection scenarios used in this study. Random selection includes random chromosome sampling. Random with inclusion of significant markers includes trait-associated markers to already selected random markers. LD selection includes selection of markers based on LD pruning; *F_ST_* scenario includes selection based on fixation index and SNP prioritization. Shannon entropy scenario includes information-based SNP selection.

#### 5.3.1 Method I - Random selection

For the first method studied, a subset of 25K, 45K, and 65K markers was randomly selected across chromosomes from the HD panel using the R software [52]. To assess sampling variation and ensure consistency of results, the random sampling scenario was repeated five times, with the average and standard deviation of the Pearson correlation coefficient assessed for the final performance of the predictions (see *“5.5 Validation strategy and performance comparison”* section). Random selection was stratified by chromosome to maintain proportional representation across the genome, i.e., larger chromosomes received more SNPs compared to smaller chromosomes. The distribution of selected markers across each scenario and chromosome is shown in S3 Table.

#### 5.3.2 Method II - Inclusion of significant and prioritized markers in the random selection approach

Using the random scenarios developed in 5.3.1, we added a total of 758 SNP markers that have been previously considered relevant for several traits in Nellore cattle. Part of these SNPs (N=298) were obtained from our gene prioritization study using 52 scientific peer-reviewed papers published in the literature for Nellore cattle [53]. The other 460 SNPs (after removing duplicates from the previously mentioned study) were identified as associated with the 16 traits currently evaluated by ABCZ (Ogunbawo et al., 2025). Consequently, the final number of SNPs in this scenario for 25K, 45K, and 65K was 25,758; 45,758; and 65,758, respectively. The distribution of selected SNPs across each scenario and chromosome is shown in S4 Table.

#### 5.3.3 Method III - Selection Based on Linkage Disequilibrium (LD)

To account for LD in SNP selection, pairwise LD was first calculated for our dataset using PLINK v1.9 [54], based on a window size of 100,000 SNPs to allow the identification of SNPs in strong LD. The descriptive statistics of the resulting LD estimates are summarized in the S5 Table. To reduce redundancy due to high LD, LD pruning was performed using sliding windows of 50 SNPs and varying *r*^2^ thresholds (i.e., 0.15, 0.2, and 0.3) for scenarios 25K, 45K, and 65K panels. The exact number of SNPs left after pruning for each scenario is shown in Table 3 and was used for further analysis. The distribution of selected SNPs across selection density and chromosome is shown in the S6 Table.

**Table 3.** Summary of SNPs retained after linkage disequilibrium (LD) pruning at different *r*^2^ thresholds and total number of SNPs in each scenario.

| $r^2$ | Selection scenario | Number of SNPs pruned | Number of SNPs left after pruning | Number of SNPs in the panel |
| --- | --- | --- | --- | --- |
| **0.15** | 25K | 407,601 | 30,049 | 30,049 |
| **0.20** | 45K | 394,345 | 43,305 | 43,305 |
| **0.30** | 65K | 365,046 | 72,604 | 72,604 |

#### 5.3.4 Method IV - Selection Based on SNP Prioritization using the Fixation Index **F_ST_** Approach

Fixation index measures genetic differentiation through changes in allele frequencies among populations [50]. In this study, to account for genetic differentiation, the global *F_ST_* estimator originally proposed by Nei (1973) was used to calculate the fixation index for all 437,650 SNPs, as well as to prioritize the SNPs to be selected [12,49,55]. Therefore, genotyped animals were divided into two subpopulations based on year of birth, with animals born between 1997 and 2009 grouped as old (N = 9,589), and animals born between 2010 and 2022 grouped as new (N = 10,000 randomly selected from 292,260). Since 292,260 animals were available in the new group, 10,000 animals were randomly selected to obtain approximately comparable subpopulation sizes. Consequently, a total of 19,589 animals were then used to calculate the F_ST_values for all SNPs, where the estimator is defined for a given locus k as follows:

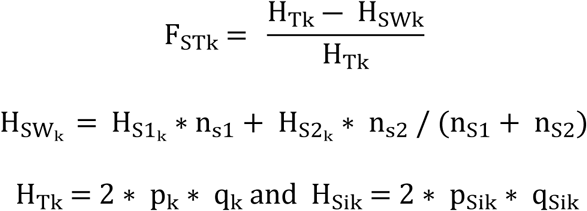

where p_Sik_ and q_Sik_ are the allele frequencies for locus k in subpopulation I of locus k, n_S1_ and n_S2_are the number of individuals in the “new” and “old” populations. The H_SWk_ is the weighted mean heterozygosity across the new and old populations, and H_Tk_ is the heterozygosity of the pooled subpopulations for locus k. For the 25K, 45K, and 65K scenarios, subsets corresponding to the top 5.7%, 10.3%, and 14.85% SNPs from the HD panel were prioritized based on their *F_ST_* thresholds corresponding to 94.3, 89.7, and 85.15, respectively, and selected SNPs were used for subsequent analyses. Descriptive statistics for the F_ST_ values of the SNPs selected in each scenario are shown in Table 4. The distribution of selected SNPs across each scenario and chromosome is shown in the S7 Table.

**Table 4.** Descriptive statistics for the F_ST_ values of the SNP selected in each scenario.

| Selection Scenario | $F_{ST}$ Cutoff | Number of SNPs in the panel |
| --- | --- | --- |
| **25K** | 0.0299 | 24,947 |
| **45K** | 0.0223 | 45,078 |
| **65K** | 0.0176 | 65,648 |

#### 5.3.5 Method V – Selection Based on Shannon entropy (Machine Learning Strategy)

This method of SNP selection is the most commonly used method in recent studies [29], as first each chromosome is divided into a fixed number of blocks, after which the Shannon Entropy is calculated at local optima and used to select SNPs [29,51]. The SNP with the highest MAF was selected, as the maximum Shannon entropy for a locus is 1 when the MAF for this locus is 0.5 [29,51]. Shannon entropy (H) is defined in information theory as the average amount of information contained in each message received [56], and is calculated as:

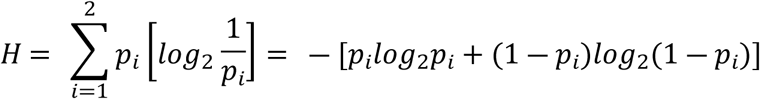

where p_i_is the minor allele frequency (MAF) of the i^th^ locus (only available for bi-allelic loci) [56]. For SNP selection, this message refers to an allele [29]. The SNP information is a function of SNP MAF and is measured by Shannon entropy as Average Shannon entropy, i.e., E-score [51]. The E-score is computed as the average of Shannon entropy across m loci for each SNP, i.e.:

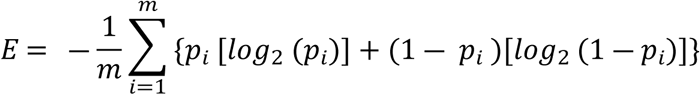

where p_i_is the MAF of SNP i, and m is the number of SNPs. For a single SNP, E is maximized and equals 1 when p_i_ = (1 ― p_i_) = 0.5 [29,51]. The average Shannon entropy for the 437,650 SNPs after the quality control was 0.7319, with a minimum of 0.2857 and a maximum of 1.0000. Of these SNPs, 205,553 had Shannon entropy greater than 0.8. These SNPs are ranked according to their Shannon entropy, after which a subset of SNPs corresponding to the top 25K, 45K, and 65K SNPs was selected. The distribution of selected SNPs across each scenario and chromosome is shown in the S8 Table.

### 5.4 Statistical Models used for the Genetic Evaluations

Genomic predictions were obtained using both the genomic best linear unbiased prediction (GBLUP) and single-step GBLUP (ssGBLUP) methods, both implemented in the BLUPF90 family of programs across selection strategies and SNP densities (including HD reference panel). The BLUPF90+ [57] was used to predict the breeding values for the linear traits (AFC, WW, YW, and MUSC), and the GIBBSF90+ software [57] was used to predict the breeding values for STAY. For STAY, a total of 10,000 iterations were generated, with the first 1,000 iterations discarded as burn-in and every 10^th^ sample retained for posterior analysis.

Raw phenotypes were used in both GBLUP and ssGBLUP rather than pseudo-phenotypes (e.g., deregressed EBVs), to maintain a consistent modeling framework and avoid introducing additional sources of variation associated with multiple-step procedures. For each trait, the GBLUP analyses were restricted to animals with both genotypic and phenotypic information. In contrast, ssGBLUP included all phenotypic records that passed quality control, and incorporated pedigree relationships from the complete pedigree containing 14,602,423 animals. To ensure a controlled comparison between methods (particularly for evaluating alternative SNP selection strategies), the genomic information used in ssGBLUP was restricted to the same trait-specific subset of genotyped animals included in the corresponding GBLUP analysis. Thus, the two methods were evaluated using the same genotyped animals and SNP panels, while ssGBLUP additionally incorporated phenotypic information from non-genotyped animals and relationships from the complete pedigree. Thus, this design was adopted to provide a balanced and interpretable comparison of the effects of SNP selection across the two methods. The number of animals in the reduced dataset used for each trait is shown in Table 2.

For both GBLUP and ssGBLUP, the genomic estimated breeding values (GEBV) for all traits except STAY were obtained by solving the mixed model equations (MME) based on the following single-trait linear mixed model:

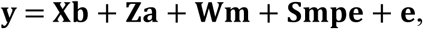

where **y** is the vector of observations for each trait; **b** is the vector of fixed effects, which included CG (herd-year-season of birth, herd-year-season of measurement, and day of measurement), age of dams class, and sex of offspring; **a** is the vector of additive genetic effect; **m** is the vector of random maternal genetic effect, **mpe** is the vector of maternal permanent environmental effects, **e** is the vector of residual effects; and **X**, **Z**, **W,** and **S** are the design matrices relating elements of **b**, **a**, **m,** and **pe** to their corresponding elements in **y**. Only the model for WW included the maternal effects.

For the GBLUP method, the expectation of random effects was 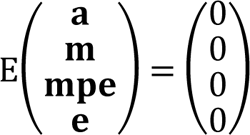, and the variance-covariance structure was defined as:

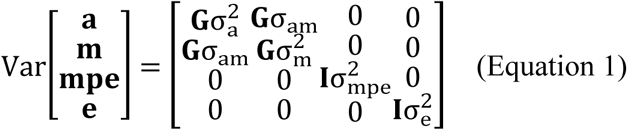

where **G** is the genomic relationship matrix created as the first method proposed in VanRaden (2008), **I** is the identity matrix, *σ^2^_a_* is the variance of the direct genetic effect, *σ^2^_m_* is the variance of the maternal genetic effect, *σ_am_* is the covariance between direct and maternal genetic effects,*σ^2^_mpe_* is the variance of the maternal permanent environmental effect, and *σ^2^_e_* is the residual variance.

For the ssGBLUP method, the **G** matrix was replaced by the hybrid relationship matrix that combines pedigree-based and genomic information (**H**) for the genotyped animals. Therefore, the inverse of the combined relationship matrix **H**^―**1**^ was created as [37,39]:

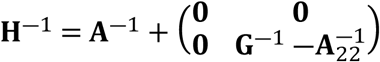

where **A**^―1^ is the inverse of the pedigree relationship matrix, A^-1^_22_ is the inverse of the pedigree relationship matrix among genotyped animals, and **G**^―1^ is the inverse of the genomic relationship matrix based on VanRaden [58].

Specifically for STAY, the following single-trait threshold model was used:

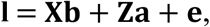

where **l** is the vector of observations (STAY) in the liability scale, assumed as l|b,a, *σ^2^_a_*, *σ^2^_e_* ~ N(Xb + Za, I*σ^2^_e_*); **b** is a vector of systematic effects (contemporary group created only based on herd-year-season of birth) assumed as b ~ N(0, I*σ^2^_b_*), where *σ^2^_b_* has a large variance to represent vague prior knowledge; **a** is a vector of additive genetic effect (random), assumed as a| *σ^2^_a_*, Σ ~ N(0, Σ*σ^2^_a_*), where **Σ** equals to **G** in GBLUP), and **H** in ssGBLUP. The *σ^2^_a_* is the additive genetic variance; **e** is the vector of residual effects assumed as e|*σ^2^_a_*, ~ N(0, I*σ^2^_e_*), where *σ^2^_e_* is the residual variance, and **I** is the identity matrix. The **X** and **Z** are the incidence matrices relating elements to the vectors of **b** and **a**, respectively.

All variance components were fixed to the estimates provided by ABCZ, which are currently used in the official evaluations of Nellore cattle performed by the association. These variance components were originally estimated using the complete datasets based solely on pedigree information. Information about the exact variance components used for each trait can be found in the S9 Table.

### 5.5 Validation strategy and performance comparison

To evaluate the performance of the 25K, 45K, and 65K SNP panels developed using the five SNP selection methods mentioned above, genomic prediction accuracy was assessed relative to the HD (reference) SNP panel comprising 437,650 SNPs (i.e., all SNPs after quality control). In addition, the SNP panel currently used by ABCZ (i.e., the 120K SNP panel [40] was also compared to the reference (HD) panel. This comparative framework allowed the assessment of whether the reduced-density panels maintain prediction accuracy compared to the current SNP panel used by the association.

The accuracy attained with each SNP panel was assessed using the Pearson correlation coefficient estimated between the GEBVs obtained from the reduced-density SNP panels (i.e., 25K, 45K, 65K, and 120K used by ABCZ) and those obtained from the reference panel (i.e., HD). High correlation coefficients (>0.95) were interpreted as indicating minimal changes in breeding decisions across selection scenarios and marker densities, suggesting that the reduced-density panels capture genetic information comparable to the reference panel. Scenarios that maintained high accuracy across the different traits and methods are preferred.

## Data Availability Statements

The raw data used in this study are available upon request to the author, Dr. Henrique Ventura. All information necessary to interpret the results and support the conclusions of this study is provided within the manuscript and its supporting information.

## Acknowledgements

The authors thank the Brazilian Association of Zebu Breeders (ABCZ) and its registered breeders for their contribution to data collection and recording. We also acknowledge the National Resource for Animal Genome Security and Productivity, currently operating as National Animal Genome Research Program (NRSP8) for supporting this research through the NRSP8 Summer Fellowship.

## Supporting Information

**S1a Table. Count of SNP markers common and unique across selection scenarios in the 25K panel**

**S1b Table. Count of SNP markers common and unique across selection scenarios in 45K panel**

**S1c Table. Count of SNP markers common and unique across selection scenarios in 65K panel**

**S2a Table. Pearson correlation across SNP density and selection scenarios using the genomic best linear unbiased prediction model for Age at first calving (AFC)**

**S2b Table. Pearson correlation across SNP densities and selection scenarios using the single step genomic best linear unbiased prediction model for Age at first calving (AFC)**

**S2c Table. Pearson correlation across SNP densities and selection scenarios using the genomic best linear unbiased prediction model for stayability.**

**S2d Table. Pearson correlation across SNP densities and Selection scenarios using the single step genomic best linear unbiased prediction model for stayability**

**S2e Table. Pearson correlation across SNP densities and selection scenarios using the genomic best linear unbiased prediction model for weaning weight**

**S2f Table. Pearson correlation across SNP densities and selection scenarios using the single step genomic best linear unbiased prediction model for weaning weight**

**S2g Table. Pearson correlation across SNP densities and selection scenarios using the genomic best linear unbiased prediction model for yearling weight**

**S2h Table. Pearson correlation across SNP densities and selection scenarios using the single step genomic best linear unbiased prediction model for yearling weight**

**S2i Table. Pearson correlation across SNP densities and selection scenarios using the genomic best linear unbiased prediction model for Muscling score**

**S2j Table. Pearson correlation across SNP densities and selection scenarios using the single step genomic best linear unbiased prediction model for Muscling score**

**S3 Table. Distribution of selected SNPs across chromosomes for the five replicates of the random selection approach at 25K, 45K, 65K SNP densities.**

**S4 Table. Distribution of selected SNPs chromosomes for inclusion of informative markers in the five replicates of random selection with informative markers at 25K, 45K, 65K SNP densities**

**S5 Table. Summary statistics of linkage disequilibrium and physical distance between SNP pairs calculated in this study.**

**S6 Table. Distribution of selected SNPs across chromosomes from the linkage disequilibrium approach at 25K, 45K, 65K SNP densities**

**S7 Table. Distribution of Selected SNPs across Chromosomes for *F_ST_* based prioritization approach at 25K, 45K, 65K densities**

**S8 Table. Distribution of Selected SNPs across Chromosomes for Shannon Entropy approach at 25K, 45K and 65K densities**

**S9 Table. Variance components used for each trait**

## Notes

### Competing Interest Statement

The authors have declared no competing interest.

## References

1. Goddard ME, Hayes BJ, Meuwissen THE. Genomic selection in livestock populations. Genet Res. 2010;92: 413–421. doi:10.1017/S0016672310000613

2. Gutierrez-Reinoso MA, Aponte PM, Garcia-Herreros M. Genomic Analysis, Progress and Future Perspectives in Dairy Cattle Selection: A Review. Animals. 2021;11: 599. doi:10.3390/ani11030599

3. Saatchi M, McClure MC, McKay SD, Rolf MM, Kim J, Decker JE, et al. Accuracies of genomic breeding values in American Angus beef cattle using K-means clustering for cross-validation. Genet Sel Evol. 2011;43: 40. doi:10.1186/1297-9686-43-40

4. Guarini AR, Lourenco DAL, Brito LF, Sargolzaei M, Baes CF, Miglior F, et al. Use of a single-step approach for integrating foreign information into national genomic evaluation in Holstein cattle. Journal of Dairy Science. 2019;102: 8175–8183. doi:10.3168/jds.2018-15819

5. Hayes BJ, Bowman PJ, Chamberlain AJ, Goddard ME. Invited review: Genomic selection in dairy cattle: Progress and challenges. Journal of Dairy Science. 2009;92: 433–443. doi:10.3168/jds.2008-1646

6. Dekkers JCM, Hospital F. The use of molecular genetics in the improvement of agricultural populations. Nat Rev Genet. 2002;3: 22–32. doi:10.1038/nrg701

7. Seidel GE. Brief introduction to whole-genome selection in cattle using single nucleotide polymorphisms. Reprod Fertil Dev. 2010;22: 138. doi:10.1071/RD09220

8. Wiggans GR, VanRaden PM, Cooper TA. The genomic evaluation system in the United States: Past, present, future. Journal of Dairy Science. 2011;94: 3202–3211. doi:10.3168/jds.2010-3866

9. Kim E-H, Kang H-C, Myung C-H, Kim J-Y, Sun D-W, Lee D-H, et al. Comparison on genomic prediction using pedigree BLUP and single step GBLUP through the Hanwoo full-sib family. Anim Biosci. 2023;36: 1327–1335. doi:10.5713/ab.22.0327

10. Gurgul A, Miksza-Cybulska A, Szmatoła T, Jasielczuk I, Piestrzyńska-Kajtoch A, Fornal A, et al. Genotyping-by-sequencing performance in selected livestock species. Genomics. 2019;111: 186–195. doi:10.1016/j.ygeno.2018.02.002

11. Meuwissen T, Hayes B, Goddard M. Genomic selection: A paradigm shift in animal breeding. Animal Frontiers. 2016;6: 6–14. doi:10.2527/af.2016-0002

12. Chang L-Y, Toghiani S, Ling A, Aggrey SE, Rekaya R. High density marker panels, SNPs prioritizing and accuracy of genomic selection. BMC Genet. 2018;19. doi:10.1186/s12863-017-0595-2

13. Wu X-L, Xu J, Feng G, Wiggans GR, Taylor JF, He J, et al. Optimal Design of Low-Density SNP Arrays for Genomic Prediction: Algorithm and Applications. PLOS ONE. 2016;11: e0161719. doi:10.1371/journal.pone.0161719

14. Matukumalli LK, Lawley CT, Schnabel RD, Taylor JF, Allan MF, Heaton MP, et al. Development and Characterization of a High Density SNP Genotyping Assay for Cattle. PLOS ONE. 2009;4: e5350. doi:10.1371/journal.pone.0005350

15. Matukumalli LK, Schroeder S, DeNise SK, Sonstegard T, Lawley CT, Georges M, et al. Analyzing LD blocks and CNV segments in cattle: novel genomic features identified using the BovineHD BeadChip. Illumina Inc: San Diego, CA. 2011.

16. Ogunbawo A, Mulim H, Campos G, Schinckel A, Oliveira H. Tailoring Genomic Selection for Bos taurus indicus: A Comprehensive Review of SNP Arrays and Reference Genomes. Genes. 2024;15: 1495. doi:10.3390/genes15121495

17. Elzo MA, Thomas MG, Martinez CA, Lamb GC, Johnson DD, Rae DO, et al. Genomic– polygenic evaluation of multibreed Angus–Brahman cattle for postweaning feed efficiency and growth using actual and imputed Illumina50k SNP genotypes. Livestock Science. 2014;159: 1–10. doi:10.1016/j.livsci.2013.11.005

18. Warburton CL, Hayes BJ. Ascertainment Bias in Cattle SNP Arrays and Implications for Multibreed Genomic Predictions. Anim Genet. 2026;57: e70081. doi:10.1002/age.70081

19. Druet T, Macleod IM, Hayes BJ. Toward genomic prediction from whole-genome sequence data: impact of sequencing design on genotype imputation and accuracy of predictions. Heredity. 2014;112: 39–47. doi:10.1038/hdy.2013.13

20. McKay SD, Schnabel RD, Murdoch BM, Matukumalli LK, Aerts J, Coppieters W, et al. Whole genome linkage disequilibrium maps in cattle. BMC Genet. 2007;8: 74. doi:10.1186/1471-2156-8-74

21. Porto-Neto LR, Kijas JW, Reverter A. The extent of linkage disequilibrium in beef cattle breeds using high-density SNP genotypes. Genet Sel Evol. 2014;46: 22. doi:10.1186/1297-9686-46-22

22. Utsunomiya YT, do Carmo AS, Carvalheiro R, Neves HH, Matos MC, Zavarez LB, et al. Genome-wide association study for birth weight in Nellore cattle points to previously described orthologous genes affecting human and bovine height. BMC Genetics. 2013;14:52. doi:10.1186/1471-2156-14-52

23. Abo-Ismail MK, Brito LF, Miller SP, Sargolzaei M, Grossi DA, Moore SS, et al. Genome-wide association studies and genomic prediction of breeding values for calving performance and body conformation traits in Holstein cattle. Genetics Selection Evolution. 2017;49: 82. doi:10.1186/s12711-017-0356-8

24. Erbe M, Hayes BJ, Matukumalli LK, Goswami S, Bowman PJ, Reich CM, et al. Improving accuracy of genomic predictions within and between dairy cattle breeds with imputed high-density single nucleotide polymorphism panels. Journal of Dairy Science. 2012;95: 4114– 4129. doi:10.3168/jds.2011-5019

25. Li W, Li W, Song Z, Gao Z, Xie K, Wang Y, et al. Marker Density and Models to Improve the Accuracy of Genomic Selection for Growth and Slaughter Traits in Meat Rabbits. Genes (Basel). 2024;15: 454. doi:10.3390/genes15040454

26. Lu D, Akanno EC, Crowley JJ, Schenkel F, Li H, De Pauw M, et al. Accuracy of genomic predictions for feed efficiency traits of beef cattle using 50K and imputed HD genotypes1. Journal of Animal Science. 2016;94: 1342–1353. doi:10.2527/jas.2015-0126

27. Pérez-Enciso M, Rincón JC, Legarra A. Sequence-vs. chip-assisted genomic selection: accurate biological information is advised. Genet Sel Evol. 2015;47: 43. doi:10.1186/s12711-015-0117-5

28. Ramos AM, Crooijmans RPMA, Affara NA, Amaral AJ, Archibald AL, Beever JE, et al. Design of a High Density SNP Genotyping Assay in the Pig Using SNPs Identified and Characterized by Next Generation Sequencing Technology. Orban L, editor. PLoS ONE. 2009;4: e6524. doi:10.1371/journal.pone.0006524

29. Wu X-L, Li H, Ferretti R, Simpson B, Walker J, Parham J, et al. A unified local objective function for optimally selecting SNPs on arrays for agricultural genomics applications. Animal Genetics. 2020;51: 306–310. doi:10.1111/age.12916

30. Abo-Ismail MK, Lansink N, Akanno E, Karisa BK, Crowley JJ, Moore SS, et al. Development and validation of a small SNP panel for feed efficiency in beef cattle1. Journal of Animal Science. 2018;96: 375–397. doi:10.1093/jas/sky020

31. Toghiani S, Aggrey SE, Rekaya R. FST-Based Marker Prioritization Within Quantitative Trait Loci Regions and Its Impact on Genomic Selection Accuracy: Insights from a Simulation Study with High-Density Marker Panels for Bovines. Genes. 2025;16: 563. doi:10.3390/genes16050563

32. Ding K, Kullo IJ. Methods for the selection of tagging SNPs: a comparison of tagging efficiency and performance. Eur J Hum Genet. 2007;15: 228–236. doi:10.1038/sj.ejhg.5201755

33. Takeuchi F, Yanai K, Morii T, Ishinaga Y, Taniguchi-Yanai K, Nagano S, et al. Linkage disequilibrium grouping of single nucleotide polymorphisms (SNPs) reflecting haplotype phylogeny for efficient selection of tag SNPs. Genetics. 2005;170: 291–304. doi:10.1534/genetics.104.038232

34. Tang NLS, Pharoah PDP, Ma SL, Easton DF. Evaluation of an algorithm of tagging SNPs selection by linkage disequilibrium. Clinical Biochemistry. 2006;39: 240–243. doi:10.1016/j.clinbiochem.2005.11.014

35. Habier D, Fernando RL, Dekkers JCM. The Impact of Genetic Relationship Information on Genome-Assisted Breeding Values. Genetics. 2007;177: 2389–2397. doi:10.1534/genetics.107.081190

36. Huang Q, Fu Y, Boerwinkle E. Comparison of strategies for selecting single nucleotide polymorphisms for case/control association studies. Human Genetics. 2003;113: 253–257. doi:10.1007/s00439-003-0965-x

37. Aguilar I, Misztal I, Johnson DL, Legarra A, Tsuruta S, Lawlor TJ. Hot topic: A unified approach to utilize phenotypic, full pedigree, and genomic information for genetic evaluation of Holstein final score. Journal of Dairy Science. 2010;93: 743–752. doi:10.3168/jds.2009-2730

38. Christensen OF, Lund MS. Genomic prediction when some animals are not genotyped. Genet Sel Evol. 2010;42: 2. doi:10.1186/1297-9686-42-2

39. Legarra A, Aguilar I, Misztal I. A relationship matrix including full pedigree and genomic information. Journal of Dairy Science. 2009;92: 4656–4663. doi:10.3168/jds.2009-2061

40. Campos G, Mulim HA, Ventura H, Souza N, Cardoso F, Oliveira HR. Genotype imputation performance in Nellore cattle across different SNP panels and software tools. Trop Anim Health Prod. 2026;58: 172. doi:10.1007/s11250-026-04914-0

41. Sargolzaei M, Chesnais JP, Schenkel FS. A new approach for efficient genotype imputation using information from relatives. BMC Genomics. 2014;15: 478. doi:10.1186/1471-2164-15-478

42. Ogunbawo AR, Hidalgo J, Mulim HA, Carrara ER, Ventura HT, Souza NO, et al. Applying the algorithm for Proven and young in GWAS Reveals high polygenicity for key traits in Nellore cattle. Front Genet. 2025;16: 1549284. doi:10.3389/fgene.2025.1549284

43. Misztal I, Tsuruta S, Strabel T, Auvray B, Druet T, Lee D. BLUPF90 and related programs (BGF90), In: CD-ROM communication, Proceedings of the 7th WORLD CONGRESS ON GENETICS APPLIED TO LIVESTOCK PRODUCTION. Montpellier. 2002;2002: 7–28.

44. Wiggans GR, Sonstegard TS, VanRaden PM, Matukumalli LK, Schnabel RD, Taylor JF, et al. Selection of single-nucleotide polymorphisms and quality of genotypes used in genomic evaluation of dairy cattle in the United States and Canada. Journal of Dairy Science. 2009;92: 3431–3436. doi:10.3168/jds.2008-1758

45. Pritchard JK, Przeworski M. Linkage Disequilibrium in Humans: Models and Data. The American Journal of Human Genetics. 2001;69: 1–14. doi:10.1086/321275

46. Stam P. The distribution of the fraction of the genome identical by descent in finite random mating populations. Genet Res. 1980;35: 131–155. doi:10.1017/S0016672300014002

47. Misztal I, Lourenco D, Legarra A. Current status of genomic evaluation. Journal of Animal Science. 2020;98: skaa101. doi:10.1093/jas/skaa101

48. Pocrnic I, Lourenco DAL, Masuda Y, Misztal I. Dimensionality of genomic information and performance of the Algorithm for Proven and Young for different livestock species. Genet Sel Evol. 2016;48: 82. doi:10.1186/s12711-016-0261-6

49. Toghiani S, Chang L-Y, Ling A, Aggrey SE, Rekaya R. Genomic differentiation as a tool for single nucleotide polymorphism prioritization for Genome wide association and phenotype prediction in livestock. Livestock Science. 2017;205: 24–30. doi:10.1016/j.livsci.2017.09.007

50. Wright S. The Genetical Structure of Populations. Annals of Eugenics. 1949;15: 323–354. doi:10.1111/j.1469-1809.1949.tb02451.x

51. Mao R, Zhou L, Wang Z, Wu J, Liu J. A Comprehensive Strategy Combining Feature Selection and Local Optimization Algorithm to Optimize the Design of Low-Density Chip for Genomic Selection. Agriculture. 2023;13: 614. doi:10.3390/agriculture13030614

52. R Core Team. R: A Language and Environment for Statistical Computing. R Foundation for Statistical Computing, Vienna, Austria. 2024. Available: https://www.R-project.org/

53. Ogunbawo AR, Mulim HA, Campos GS, Oliveira HR. Genetic Foundations of Nellore Traits: A Gene Prioritization and Functional Analyses of Genome-Wide Association Study Results. Genes. 2024;15: 1131. doi:10.3390/genes15091131

54. Purcell S, Neale B, Todd-Brown K, Thomas L, Ferreira MAR, Bender D, et al. PLINK: a tool set for whole-genome association and population-based linkage analyses. Am J Hum Genet. 2007;81: 559–575. doi:10.1086/519795

55. Nei M. Analysis of Gene Diversity in Subdivided Populations. Proceedings of the National Academy of Sciences. 1973;70: 3321–3323. doi:10.1073/pnas.70.12.3321

56. Shannon CE. A mathematical theory of communication. The Bell System Technical Journal. 1948;27: 379–423. doi:10.1002/j.1538-7305.1948.tb01338.x

57. Misztal I, Lourenco D, Aguilar I, Legarra A. Manual for BLUPF90 family of programs. 2014. Available: http://nce.ads.uga.edu/wiki/lib/exe/fetch.php?media=blupf90_all.pdf

58. VanRaden PM. Efficient Methods to Compute Genomic Predictions. Journal of Dairy Science. 2008;91: 4414–4423. doi:10.3168/jds.2007-0980

